# Estimating partial body ionizing radiation exposure by automated cytogenetic biodosimetry

**DOI:** 10.1101/2020.09.01.278200

**Authors:** Ben C. Shirley, Joan H.M. Knoll, Jayne Moquet, Elizabeth Ainsbury, Pham Ngoc Duy, Farrah Norton, Ruth C. Wilkins, Peter K. Rogan

## Abstract

**Purpose:** Inhomogeneous exposures to ionizing radiation can be detected and quantified with the Dicentric Chromosome Assay (DCA) of metaphase cells. Complete automation of interpretation of the DCA for whole body irradiation has significantly improved throughput without compromising accuracy, however low levels of residual false positive dicentric chromosomes (DCs) have confounded its application for partial body exposure determination.

**Materials and Methods:** We describe a method of estimating and correcting for false positive DCs in digitally processed images of metaphase cells. Nearly all DCs detected in unirradiated calibration samples are introduced by digital image processing. DC frequencies of irradiated calibration samples and those exposed to unknown radiation levels are corrected subtracting this false positive fraction from each. In partial body exposures, the fraction of cells exposed, and radiation dose can be quantified after applying this modification of the contaminated Poisson method.

**Results:** Dose estimates of three partially irradiated samples diverged 0.2 to 2.5 Gy from physical doses and irradiated cell fractions deviated by 2.3-15.8% from the known levels. Synthetic partial body samples comprised of unirradiated and 3 Gy samples from 4 laboratories were correctly discriminated as inhomogeneous by multiple criteria. Root mean squared errors of these dose estimates ranged from 0.52 to 1.14 Gy^2^ and from 8.1 to 33.3%^2^ for the fraction of cells irradiated.

**Conclusions:** Automated DCA can differentiate whole-from partial-body radiation exposures and provides timely quantification of estimated whole-body equivalent dose.

**Biographical Note:** Ben Shirley M.Sc. is Chief Software Architect, CytoGnomix Inc. Canada; Joan Knoll Ph.D. Dipl.ABMGG, FCCMG is Professor in Pathology and Laboratory Medicine, Schulich School of Medicine and Dentistry, University of Western Ontario, Canada and cofounder, CytoGnomix Inc.; Jayne Moquet Ph.D. is Principal Radiation Protection Scientist in the Cytogenetics Group, Public Health England; Elizabeth Ainsbury Ph.D. is Head, Cytogenetics Group and the Chromosome Dosimetry Service, Public Health England; Pham Ngoc Duy M.Sc. is deputy director of Biotechnology Center, Dalat Nuclear Research Institute, Vietnam; Farrah Norton M.Sc.is Research Scientist and Lead of the Biodosimetry emergency response and research capability at Canadian Nuclear Laboratories; Ruth Wilkins, Ph.D. is Research Scientist and Chief of the Ionizing Radiation Health Sciences Division at Health Canada, Ontario, Canada; and Peter K. Rogan Ph.D. is Professor of Biochemistry and Oncology, Schulich School of Medicine and Dentistry, University of Western Ontario, Canada, and President, CytoGnomix Inc.

## Introduction

Accurate biological doses received by individuals exposed to ionizing radiation must be determined in order to effectively diagnose and treat victims. The dicentric chromosome assay (DCA) is the gold standard biological dose assessment method and is endorsed by the International Atomic Energy Agency (IAEA), the World Health Organization, and the Pan American Health Organization. Dicentric chromosome (DC) aberrations are biomarkers of radiation exposure and the IAEA recommends a sufficient count of either images examined or DCs encountered for accurate assessment of biological dose. Low linear energy transfer (LET) generates chromosome breaks that can be mis-repaired as DCs, which exhibit a Poisson distribution in cells. However, if radiation exposure is inhomogeneous (partial body), the portion of exposed cells expected to conform to a Poisson distribution of DCs must be determined prior to estimating absorbed dose (ISO 19238 2004; ISO 21243 2008; International Atomic Energy Agency 2011).

Traditionally, interpreting the DCA is a painstaking process which requires significant training to perform. Following extensive laboratory processing (Oestreicher et al. 2017), the operator examines metaphase images, excludes those of poor quality, documents DCs in each image, then determines the overall frequency of DCs. The frequency of DCs per cell is related to absorbed radiation dose (in Gray [Gy]). The DCA has been shown to be accurate for the 0-5 Gy range of exposures by fitting DC frequencies of known dose to a linear-quadratic calibration curve. The absorbed dose of samples of unknown exposure is inferred from the calibration curve based on DC frequency. For accurate dose assessment, detection of at least 100 DCs at higher doses is recommended. However, at low dose or partial body exposures in which DCs are much less frequent, scoring of several thousand images is necessary for accurate dose estimation (International Atomic Energy Agency 2011) (though scoring of fewer cells is recommended as a first step to handle large numbers of samples for rapid ‘triage’ in emergency response (Oestreicher et al. 2017)).

Automated approaches have been sought after to improve the throughput of the DCA, especially for large scale testing (Maznyk et al. 2012). Semi-automated detection of DCs still requires manual image selection and verification of candidate DCs (Schunck et al. 2004). The Automated Dicentric Chromosome Identifier and Dose Estimator (ADCI) software completely automates DC detection and estimates biological radiation dose (Rogan et al. 2016). Suboptimal metaphase images are removed (Liu et al. 2017), chromosomes within remaining images are classified, which are then further discriminated as either normal or DC. ADCI generates calibration curves and estimates exposure levels of samples of uncertain dose. ADCI can process a sample of 500 metaphase images and estimate dose in ~3-5 minutes using a multicore desktop computer system (Intel i7-6700HQ, 16Gb RAM) equipped with a graphics processing unit (GPU; Nvidia^®^ GTX 960M or RTX 2070) (Li et al. 2019). This benchmark estimate is equivalent to ~1.7 images per second, or ~6000 images per hour.

Image selection models which eliminate and/or rank images are a prerequisite for accurate automated dose estimation. The models are optimized to filter out suboptimal chromosome morphology and control for preparation differences that are often variable between laboratories. Application of these models can significantly reduce misclassification of DCs and increase the accuracy of DC frequencies (Shirley et al. 2017).

Nevertheless, residual False Positive (FP) DCs, that is, monocentric chromosomes incorrectly classified as DCs, produce inflated dose estimates, especially in samples exposed to low levels of radiation. A previously published FP method removes FP DCs flagged by ADCI by applying filters designed to detect morphological subclasses of FPs (Liu et al. 2017). These chromosomes are reclassified as normal, monocentric chromosomes and can be visualized in ADCI in the built-in Metaphase image viewer. While 55% of FPs on average are eliminated using this method, some FPs remain after filtering. The impact of the residual FPs is minimal when both calibration and test samples are processed using the same algorithm, resulting in the equivalent levels of FP misclassification in all images, regardless of source. This effectively mitigates their effect on dose estimation (Li et al. 2019). Dose estimation accuracy is therefore unaffected, and results fulfill IAEA criteria for triage biodosimetry.

Heterogeneous, partial body exposure is prevalent in cases of accidental radiation exposure (Prasanna et al. 2010). Partially irradiated samples deviate from the expected Poisson distribution, as the unirradiated portion of cells inflates the percentage of cells lacking DCs. This deviation must be considered to avoid underestimating exposures. The impact of FP DCs on dose estimates of partially irradiated samples was not predictable and affected the accuracy of some estimates, especially at low dose exposures. We describe a framework for automated estimation of partially irradiated samples using ADCI, which effectively corrects DC counts of FPs resulting from image segmentation and machine-learning based misclassification (Figure 1).

**Figure 1.**
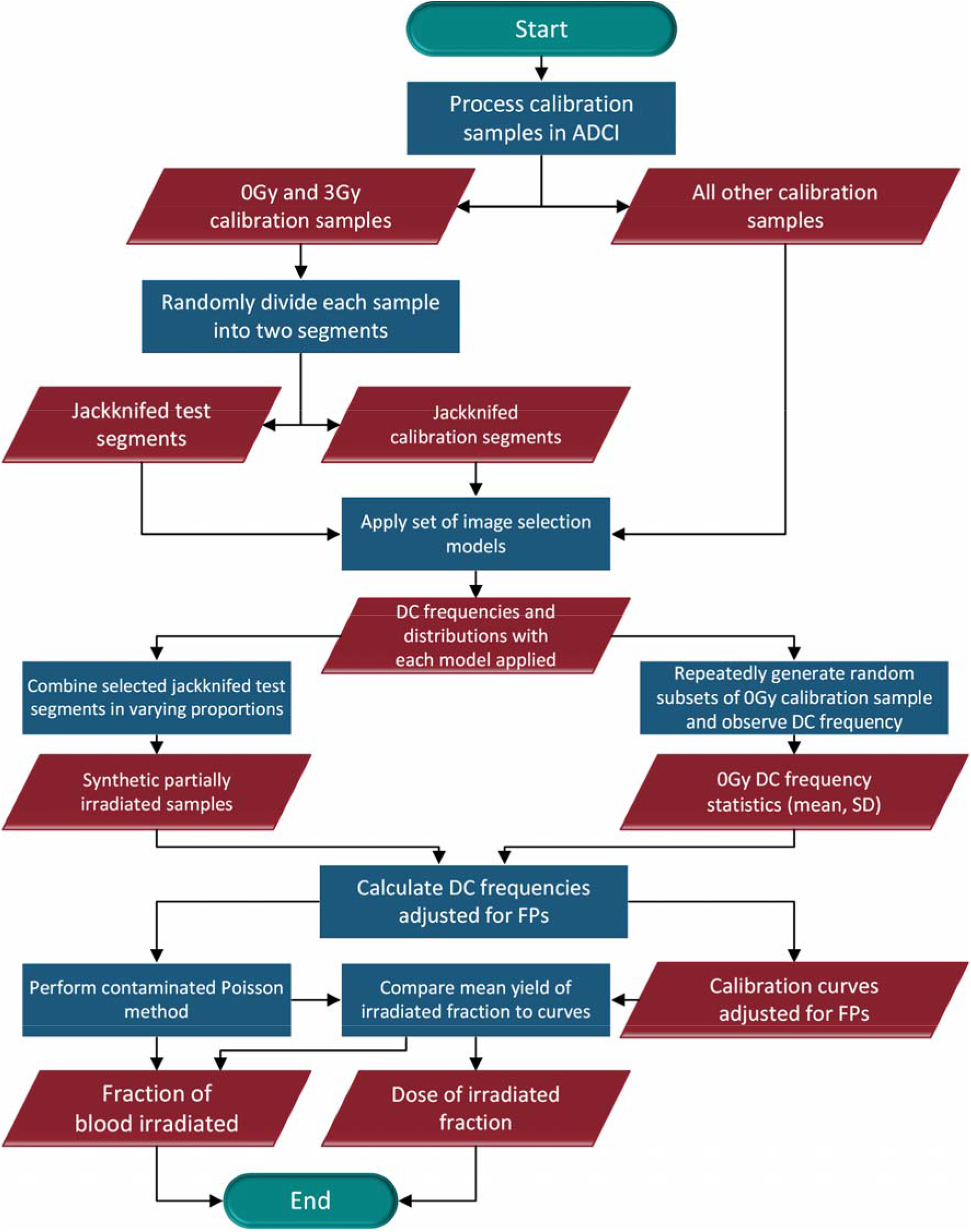
Flowchart of major steps taken to synthesize partially irradiated samples, perform dose estimation, and predict fraction of cells irradiated. Rounded shapes denote start and end points, rectangular shapes represent operations which must be performed, slanted parallelograms represent datasets. The flowchart presents the steps necessary to analyze samples originating from a single laboratory, all steps were repeated for each laboratory.

## Methods

### Sample preparation and image capture

Samples were irradiated by biodosimetry laboratories at Health Canada (HC), Canadian Nuclear Laboratories (CNL), Public Health England (PHE), and Dalat Nuclear Research Institute (DNRI) using established protocols (International Atomic Energy Agency 2011; Oestreicher et al. 2017; Pham et al. 2019). HC irradiated samples using 250 kVp X-rays (X-RAD-320 (Precision X-ray, North Branford, CT)) at a dose rate of 0.8 Gy/min, CNL used a ^137^Cs GammaCell40 (Atomic Energy of Canada Ltd, Ottawa, ON) at a dose rate of ~4.5 rad/sec, DNRI used 200 kVp X-rays (Radioflex-200EGM (Rigaku, Japan)) at a dose rate of 0.497 Gy/min. Samples obtained from PHE were irradiated *ex vivo* in a water phantom at 37 °C to ^60^Co gamma rays, with a dose rate of 0.27 Gy/min, at the University of Ghent irradiation facility. Dosimetry was performed with a NE2571 Farmer ionization chamber (Thermo Electron, UK) calibrated in terms of air kerma using the IAEA TRS-277 code of practice. To simulate partial body irradiations, irradiated blood samples were mixed with sham-irradiated control blood from the same donor in a ratio of 1:1 and sent to PHE, at room temperature, for processing using standard techniques (International Atomic Energy Agency 2011).

All laboratories captured images of metaphase cells utilizing a Metafer slide scanning platform (Metasystems, Newton, MA). HC scanned slides on a Zeiss AxioImager.Z2 microscope connected through a CoolCube 1 CCD camera using Metafer4 v3.10.7 software. CNL scanned slides on a Zeiss AxioImager.Z2 microscope equipped with a CoolCube 1 CCD camera using Metafer4 v3.11.8 software. PHE scanned slides on a Zeiss AxioImager.M1 microscope and CoolCube 1 CCD camera using Metafer4 v3.9.10 software. PHE manually selected images that appeared to contain approximately 46 chromosomes of good morphology that were reasonably well spread from the low magnification (10X) scan image gallery. DNRI scanned slides on an AxioImager.Z2 microscope with CCD camera using Metafer4 v3.10 software. DNRI further selected images based on the following criteria: metaphase cells at first mitotic division post-irradiation, with 46 chromosomes that are nonoverlapping, well spread with chromatids separated. HC, DNRI, and PHE utilized MSearch to eliminate images lacking metaphase cells. CNL used MSearch to capture all unsorted images automatically without applying any selection criteria; ADCI was used to eliminate those which did not contain metaphase cells. Images were exported as TIFF files.

### Sample transfer and image processing

Calibration samples of known dose ranging from 0-5Gy (0-4.5Gy for PHE, 0-4Gy for HC) were obtained from each laboratory. All test samples were derived from whole-body (WB) exposures, except for PHE, which provided four WB and three partial-body (PB) irradiated samples. Except for HC and CNL (Li et al. 2019), transfer of metaphase image data was performed via secured internet connection using Synology Cloud Station software to a centralized Network Attached Storage device at the University of Western Ontario. Results for each laboratory were separated, and images were grouped by sample dose. Transfers took 12 to 24 hours on average, depending on image count and internet connection speed. To assess transfer success, file counts were matched to the expected number of images, and random images were opened to assess potential data corruption.

ADCI software was used to examine metaphases using image processing, image segmentation, and machine learning methods (Rogan et al. 2016; Li et al. 2016; Shirley et al. 2017; Liu et al. 2017; Li et al. 2019). This process removes irrelevant objects/debris, locates candidate centromeres, and discriminates dicentric from monocentric chromosomes. A chromosome-level filtering algorithm removed the majority of false positive (FP) DCs.

Image selection models in ADCI first exclude suboptimal images based on filter criteria described below, then optionally rank and select a specified number of remaining images. The IAEA recommends examination of 500-1000 images when estimating dose (International Atomic Energy Agency 2011). A target minimum of 1000 images was set due to the overdispersed distribution of DCs found in partially irradiated samples. Therefore, images in jackknifed samples were selected using models which select the 500 top ranked images. When combined, two jackknifed samples will produce a synthetic sample containing at least 1000 images. However, partially irradiated samples obtained from PHE contained only 899 or 900 images. For these samples, an image selection model which selected the optimal 750 images was used.

Image selection models comprising criteria to filter out suboptimal metaphase cell images were chosen for each laboratory based on highest dose estimation accuracy for homogeneous radiation exposures (Li et al. 2019; Rogan et al. 2019). These filters include: I) Length-Width Ratio: which removes cells with excessively long or thin chromosomes, II) Centromere Candidate Density: which removes cells with chromosomes exhibiting high densities of centromere candidates, III) Finite Difference: which removes cells containing excessive numbers of objects with smooth contours, such as nuclei or micronuclei, IV) Object Count: which eliminates images with excessive or insufficient chromosome counts due to excessive sister chromatid separation, debris or multiple metaphase cells, V) Segmented Object Count: which removes images with excessive or sparse numbers of segmented objects, and VI) Classified Object Ratio: which removes cells with an insufficient fraction of objects that are recognized as chromosomes. Images can also be ranked according to the area distributions of chromosome objects based on the degree to which they adhere to the natural distribution of chromosome lengths in a normal karyotype (Group Bin method) (Liu et al. 2017). To ensure adequate image counts for partial body dose estimation, jackknifed samples from CNL, PHE, and DNRI processed with previous models (Li et al. 2019), that selected fewer than 500 images, were re-evaluated with models requiring at least this number of metaphase cell images. ADCI can also generate optimal image selection models programmatically by exhaustively searching all models within specified ranges of filtering thresholds (Li et al. 2019). Automated Image Selection Model 48735 applied filters II, IV, VI, and selected the 500 top images ranked by Group Bin Distance was applied to HC synthetic samples and was used to minimize HC calibration curve fit residuals. The C_B500 and C_B750 models differ only in the number of images selected and apply image Filters IIII to each set of images then select 500 or 750 remaining images ranked according to Group Bin Distance. Model C_B500 was applied to CNL, PHE, and DNRI synthetic samples. Model C_B750 was applied to partially irradiated samples obtained from PHE.

### Adjustment of DC frequencies to correct for misassigned FP dicentric chromosomes

FP DCs flagged by ADCI have been minimized using morphological image filtering (Liu et al. 2017), however not completely eliminated. We correct the overall DC frequencies by subtracting the estimated residual FPs after morphologic filtering in each sample. The residual FPs were previously reported to be distributed uniformly across doses from 0-5 Gy, because they originate from limitations in the algorithm used to detect them (Liu et al. 2017). Based on previous studies that have determined DC frequencies in unirradiated normal tissues, we assume that nearly all DCs detected in 0 Gy calibration samples are FPs. The DC frequencies of irradiated calibration samples are processed by subtracting the 0 Gy FP fraction from each. The corrected DC frequencies of irradiated calibration samples are used to generate a calibration curve adjusted for FPs identified by ADCI. The FP DC count in the 0 Gy calibration sample (0*Gy FP*) is corrected to eliminate DCs above baseline rates, which have been shown to occur at a frequency of 0.00078 in unirradiated cells (Lloyd et al. 1980):

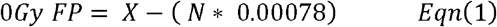

where *X* is the number of observed DCs and *N* is the total cell count. True positive (TP) DCs present in each >0 Gy calibration sample can be determined using the following equation:

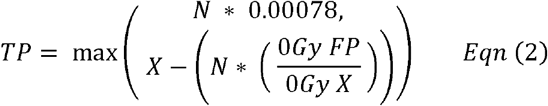

where *X* is the number of observed DCs in the >0 Gy calibration sample and 0*Gy X* is the number of observed DCs in the 0 Gy sample. This equation ensures the TP DC count in all >0 Gy calibration samples cannot fall below the expected DC rate in an unirradiated sample. Finally, the adjusted DC frequency of each sample is calculated using TP count in place of *X* when dividing by *N*.

Dolphin (World Health Organization & International Atomic Energy Agency 1969) introduced the contaminated Poisson method for estimating partial body exposures. The IAEA manual defines key equations as follows:

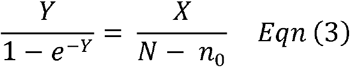

where *Y* is the mean yield of DCs in the irradiated fraction, *e-^Y^* represents cell count with no DCs in the irradiated fraction, *X* is the number of observed DCs, *N* is the total cell count, and *n_0_* is the total cell count which contain no DCs. At this point, *Y* can be compared to a calibration curve, resulting in an estimated dose of the irradiated fraction (*D*). In order to determine the fractions of cells irradiated, it is necessary to use Eq. (4) to estimate the fraction of irradiated cells which reach metaphase (*p*) after taking into account interphase death and mitotic decay:

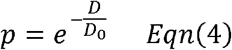

where *D_0_* is the dose at which 37% of irradiated cells survive. The value of *D_0_* is dependent on the radiation source and can vary from study to study. The *D_0_* value of 3.8 Gy (Barquinero et al. 1997) was assigned for HC and DNRI samples exposed to X-rays and 3.5 Gy (Matsubara et al. 1974) for CNL and PHE samples exposed to ^60^Co gamma rays. The estimated fraction of cells irradiated (*F*) can be determined as follows:

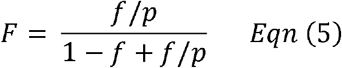

where *f* is the fraction of observed cells which were irradiated.

The method assumes that the DC frequency at 0 Gy is accurate and consistent across both calibration and test samples, which has been previously demonstrated in multiple studies (Lloyd et al. 1980; International Atomic Energy Agency 2001). We developed software to randomly select a specified number of images from any sample processed by ADCI. For each laboratory, we used this script to generate 500 randomly selected subsets of samples, each subset containing half the number of images present in the original 0 Gy sample, or 500 images, whichever was greater. Image selection models were then applied to identify the optimal metaphase cell images in each subset. The image selection models applied consisted of either a minimal model, excluding only those images in which ADCI cannot locate a metaphase cell, or an optimal model previously determined to select images fulfilling IAEA triage dose estimation criteria when estimating dose for homogeneous radiation exposures. We examined the DC frequency of each image selected subset and determined variance, standard deviation (SD), coefficient of variation, and the maximum and minimum values of DC frequency among the subsets (Supplementary Table 1). For each laboratory, a histogram was created to indicate the distribution of DC frequencies across the randomly generated samples. Histogram binning and calculation of Gaussian curve fit overlay using nonlinear regression were performed using GraphPad Prism version 6.07.

Adjusted DC frequency values were manually entered into ADCI to generate calibration curves adjusted for FPs. Best fit linear quadratic coefficients were determined using the maximum likelihood method (Papworth 1975). Resultant curves are highly similar to unadjusted curves in shape, shifted to a lower set of DC frequencies. For each laboratory, an additional curve was generated by reducing the observed 0 Gy calibration sample DC frequency by two SD. In practice, this was accomplished by reducing *X* in Eq. (1) in accordance with a reduction in 0 Gy DC frequency by two SD, then performing equations 2 through 5 as normal. DCs are expected to be very infrequent in 0 Gy samples, so those observed are highly likely to be FPs introduced during image processing. The 0 Gy calibration sample DC frequency reduced by two SD is a conservative estimate of the minimum DC frequency which could reasonably be found in the sample. By removing this reduced number of FPs in test samples, we can ensure that DC counts in test samples after FP adjustment are non-negative.

### Creation of synthetic, partially irradiated samples from mixtures of unirradiated and radiation-exposed metaphase cell images

One half of the metaphase images from the 0 Gy and 3 Gy calibration samples from the same laboratory were randomly selected to simulate new jackknifed samples for use as partial body radiation test samples. The remaining unselected images were used for generation of calibration curves.

Partially irradiated samples were constructed from the jackknifed 0 Gy and 3 Gy test image pool of each laboratory by varying the proportions of irradiated fraction in the synthetic sample (with the 3 Gy fraction representing either 9.1%, 16.7%, 25%, 33.3%, 50%, or 66.7% of the total). Image selection models were applied within ADCI before these samples were combined, allowing a specified number of top-ranking images from each jackknifed sample to be combined. To achieve the proportions listed above, top images from necessary samples were added multiple times to the synthetic sample. For example, to construct a sample with 33.3% of cells irradiated, top images from a 0 Gy sample appear twice in the constructed sample while top images from a 3 Gy sample appear once. Samples were constructed in this manner using a software script which directly interprets processed images, alleviating the need to reprocess each constructed sample using ADCI.

To control for potential sample bias during the jackknife procedure, a second set of synthetic samples was created by swapping calibration and test portions of jackknifed samples, then repeating the steps described above. We define the first set of synthetic samples as “synthetic sample set A” and the samples created after swapping calibration and test portions as “synthetic sample set B”.

### Dose estimation of samples with heterogeneous exposures

DC frequencies of calibration samples were determined after application of an image selection model. The same image selection model is then applied to test samples. If a sample has been partially irradiated, estimated dose of irradiated fraction and estimated fraction of blood irradiated are determined by utilizing the contaminated Poisson method (IAEA 2011). Calibration curve generation and dose estimation of test samples was repeated using either the optimum preset image selection model or automatically generated model for each laboratory.

Formulae to determine the mean DC yield in the irradiated fraction and the actual fraction of cells irradiated were implemented as C++ software (associated source code, spreadsheet, and example data available in the Zenodo archive doi:10.5281/zenodo.3908607) (World Health Organization & International Atomic Energy Agency 1969; International Atomic Energy Agency 2011). The solution to the mean yield of DCs in the irradiated fraction (*Y*) equation was approximated using bisection (Boost library version 1.62).

The distribution of DCs across all metaphase cells in a sample is required for the contaminated Poisson method. The DC distribution can be obtained through the console in ADCI which displays categories, i.e. the number of metaphase cell images in a sample containing 0 DCs, 1 DC, and so on. The method corrects for FPs introduced during cell image analysis, which can distort estimates of partial body dose and fraction of cells irradiated. FPs are removed from each category in the distribution in a uniform manner to maintain the overall percentage of images in each bin. Since all relevant terms in formulae necessary to perform the contaminated Poisson method can be represented as decimal numbers, the DC counts can be directly adjusted without having to round them to the nearest integer.

For each test sample, *Y* was compared to calibration curves adjusted for FPs resulting in estimated partial body doses (*D*). Confidence intervals for *D* were computed by substituting *Y* + SD and *Y* - SD in place of *Y* when comparing *Y* to the calibration curve. Confidence intervals for *F* were computed by substituting upper and lower confidence interval values for *D* in place of *D* and then repeating equations 4 and 5.

### Detection of partially irradiated samples

The *u* test, a measure of fit to a Poisson distribution, indicates overdispersion (*u*>1.96) and underdispersion (*u*<-1.96) when present (Rao & Chakravarti 1956; Savage 1970). Irradiation of a fraction of cells is often evident from overdispersion of DCs in samples that significantly differ from the Poisson distribution. The *u* values presented here were determined from the DC distribution in samples after application of an image selection model, but before adjustment for FPs.

Besides the *u* test, laboratories also compare estimated whole body and partial body doses (International Atomic Energy Agency 2011). If the two dose estimates differ significantly, this provides further evidence a sample may have been received inhomogeneous exposure. *u* values and partial body radiation dose estimates were generated for exercise samples known to be homogeneously irradiated from CNL, HC, PHE, and DNRI (9, 6, 4, and 3 samples respectively). Three partially irradiated samples from PHE (labeled E, F, and G) were also examined.

## Results

When inhomogeneous radiation exposure is suspected, samples must be classified as either whole body or partially irradiated in order to generate accurate dose estimates. Quantification of partial body radiation by either the contaminated Poisson or Qdr methods (International Atomic Energy Agency 2011) is then used to determine the values of *D* and *F*, respectively. Our previous efforts derived image filtering methods that eliminated the majority of FPs found by ADCI, while maintaining all of the true DCs detected in these samples (Rogan et al. 2016). A modified contaminated Poisson approach for partial body radiation assessment adjusted the observed DC counts to correct for residual FPs. Correction of DC counts used unirradiated cells in which DCs are rare. Estimates of both dose (*D*), and to a greater extent, the fraction of exposed cells (*F*) significantly improved in nearly all cases after DC counts were corrected.

### Dicentric distribution adherence to Poisson distribution

Adherence to the expected Poisson distribution was evaluated based on the *u* value for the seven test samples obtained from PHE, including three partially irradiated samples (Table 1a). PHE samples A-D were included to act as an additional set of homogeneously exposed controls. Unmodified and jackknifed 0 and 3 Gy calibration samples from each laboratory were also examined (Table 1b). Bolded *u* values denote a correct classification, corresponding to *u*≤1.96 for homogeneously exposed samples and *u*>1.96 for partially irradiated samples.

**Table 1a.** Computed *u* values of PHE test samples

|  | Homogeneous radiation exposure |  |  |  | Partially irradiated |  |  |
| --- | --- | --- | --- | --- | --- | --- | --- |
|  | PHE_A | PHE_B | PHE_C | PHE_D | PHE_E | PHE_F | PHE_G |
| Minimal | **1.76** | **0.18** | 2.48 | 2.08 | **5.41** | 1.95 | **4.23** |
| Optimal | **0.23** | **0.16** | **0.38** | 2.38 | **6.15** | 0.60 | **2.28** |

**Table 1b.** Computed *u* values of unmodified and jackknifed 0 Gy and 3 Gy calibration samples

| Dose | Sample type | Lab: |  | CNL |  | HC |  | PHE |  | DNRI |  |
| --- | --- | --- | --- | --- | --- | --- | --- | --- | --- | --- | --- |
|  |  |  |  | M* | O^ | M | O | M | O | M | O |
| 0 Gy | Unmodified Calibration |  |  | 92.39 | 2.10 | 14.20 | 2.55 | 7.60 | **-0.43** | 6.43 | **0.70** |
|  | Jackknifed Calibration |  |  | 67.54 | 6.18 | 8.54 | 3.49 | 3.82 | **-0.36** | 6.38 | 6.87 |
|  | Jackknifed Test |  |  | 63.13 | **0.60** | 11.60 | **-0.28** | 7.14 | **-0.49** | 2.93 | **-0.50** |
| 3 Gy | Unmodified Calibration |  |  | 51.71 | **1.72** | 6.69 | **0.10** | 1.50 | **0.49** | 2.37 | **0.68** |
|  | Jackknifed Calibration |  |  | 39.11 | -<br>**0.80** | 4.95 | **0.54** | 2.03 | **1.20** | 2.32 | **1.42** |
|  | Jackknifed Test |  |  | 33.96 | 2.94 | 4.52 | **-0.28** | 0.13 | **0.73** | 0.95 | **0.73** |

\* M denotes usage of a minimal image selection model which in almost all cases selects all images
^ O denotes usage of an optimal image selection model

These results show that an effective image selection model is frequently required for samples of homogeneous exposure in order to obtain the expected result of *u*≤ 1.96. Minimal image selection produced 21/24 unmodified and jackknifed calibration samples that were incorrectly classified as overdispersed. After application of the optimal image selection model, 18/24 samples were correctly classified as homogeneously exposed and 5/6 misclassifications were made as a result of excess DCs in unirradiated (0 Gy) samples. Examination of PHE samples follows a similar trend with 3/4 homogenously exposed samples correctly classified, and the only misclassification occurred with the unirradiated sample, PHE_D. Partially irradiated samples PHE_E (4 Gy, 50% fraction) and PHE_G (6 Gy, 50% fraction) were correctly classified as overdispersed. PHE_F (2 Gy, 50% fraction), the lowest dose sample, was not recognized as a partial body exposure.

### Variation within samples

Subsets of jackknifed calibration data from 0 Gy samples from each laboratory were randomly sampled (n=500) to obtain a set of distributions of DC frequencies (Figure 2, inset). The degree to which variation exists within an unirradiated sample quantified the expected DC frequency range. In general, the standard deviation of DC frequency is inversely related to sample image count, however image selection models specifying a maximum image count have a lower standard deviation than the same image count selected at random.

**Figure 2.**
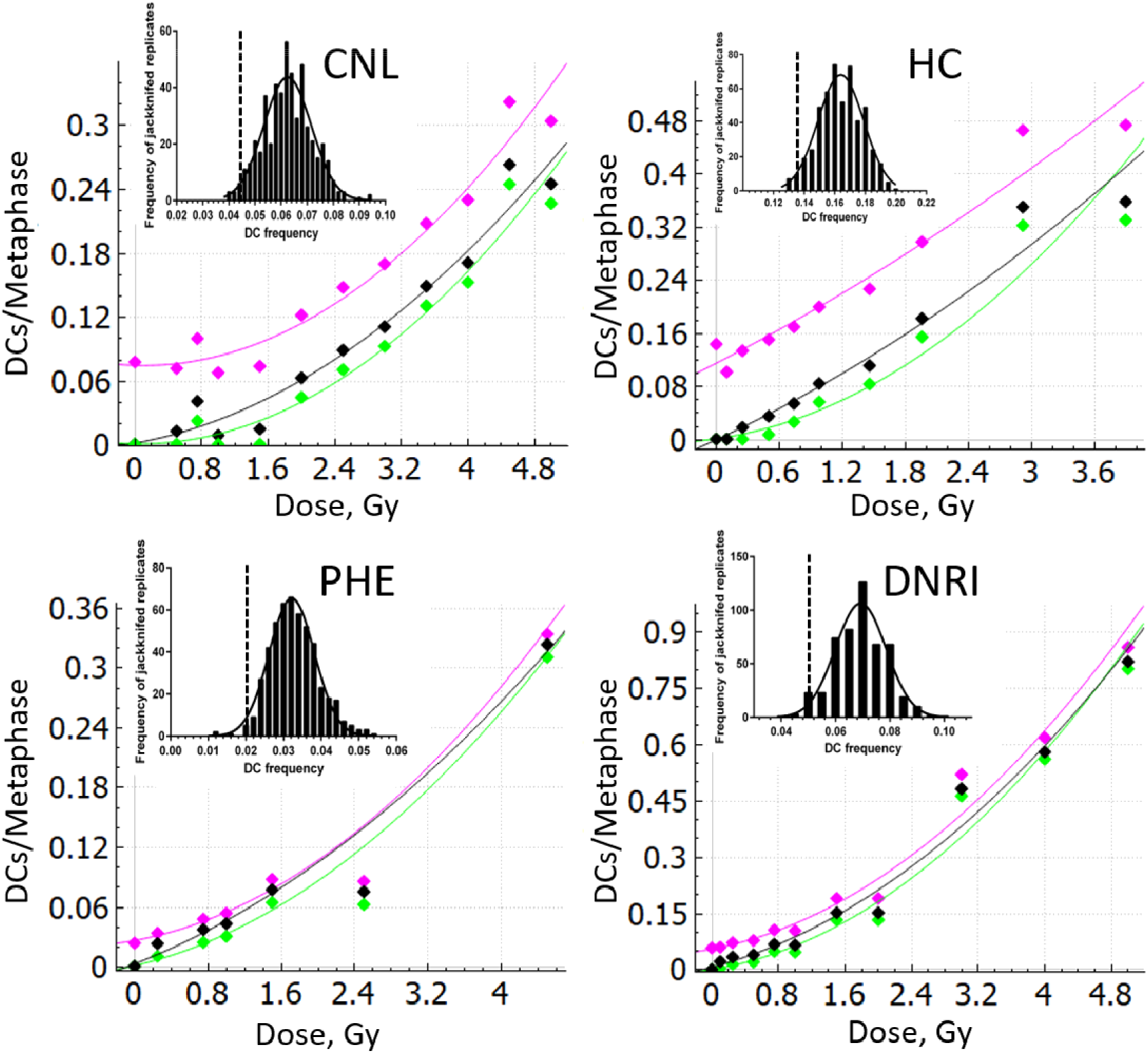
Calibration curves generated by ADCI before FP adjustment [purple colour in the digital version of this study], after FP adjustment [green], and after both FP adjustment and a DC frequency reduction by 2 standard deviations from the mean of frequency from randomly selected cell images of a series of subsets [black]. Insets within each plot contains a histogram presenting the distribution of DC frequency across 500 sample subsets, each generated by randomly selecting half of the images from the corresponding 0 Gy jackknifed calibration sample. A Gaussian curve was fit to the histogram values. The dashed line denotes 2*SD below the mean.

Adjusted calibration curves based on calibration samples with an image selection model applied were generated for all laboratories (Figure 2). Coefficients for curve fitting and *R*^2^ values for these curves are provided in Supplementary Table 1. In all cases the y intercept is decreased (as expected) by the FP adjustment. In all cases, the quadratic component of the curve is decreased by the FP adjustment, but the decrease is even greater by applying the correction to prevent negative DC counts. The extent of the correction is minimal for DNRI and PHE. Since the quadratic component has a larger impact at high radiation exposures, applying this correction to calibration curves will distort high dose exposure estimates to a greater degree in samples from HC and CNL than the other laboratories. In contrast with DNRI and PHE, images from these laboratories were not manually processed or reviewed prior to analysis using ADCI (Li et al. 2019). Based on previously published studies (Lloyd et al. 1980), all 0 Gy jackknifed samples were specified to exhibit DC frequencies of 0.00078 after FP adjustment. DC frequencies for other doses were reduced proportionately by the frequency of FP DCs predicted to exist in the 0 Gy sample before adjustment. This relationship is illustrated by observing the curve generated after FP adjustment (green in Figure 2), all DC frequencies have been uniformly reduced. DC frequencies in >0 Gy calibration samples adjusted for FPs are higher in curves generated with 0 Gy DC frequency decreased by 2*SD, and lower in curves generated with 0 Gy DC frequency with no additional standard reduction based on standard deviation. Dose estimates of partially irradiated samples utilized the calibration curve adjusted for FPs, which was based on the corrected DC frequency of the 0 Gy calibration sample, less 2*SD.

### Assessment of partially irradiated samples

PHE test samples E, F, and G consist of equal proportions of irradiated and unirradiated cell samples, which is typical of partial body irradiation. The estimated dose of the irradiated fraction and fraction of cells irradiated are presented in Table 2. The overall Root Mean Squared Errors (RMSE) for the PHE partial body samples of the dose estimate is 1.59 Gy^2^ and is 9.33%^2^ for the fraction of cells irradiated. The dose-estimated RMSE for these samples is comparable to whole-body estimated dose errors from unselected cells (Li et al. 2019).

**Table 2.** Estimated dose and predicted fraction of cells irradiated in PHE samples with heterogeneous radiation exposure

| Sample | Dose (Gy) | Estimated total body irradiation exposure (Gy) |  | Estimated exposure to irradiated fraction (Gy) |  | Predicted fraction of blood irradiated (%) |  |
| --- | --- | --- | --- | --- | --- | --- | --- |
|  |  | M* | O^ | M | O | M | O |
| PHE_E | 4.0 | 1.0 [0.3,1.7] | 1.6 [0.9,2.4] | 5.6 [3.7,7.4] | 5.2 [4.0,6.2] | 47.3 [26.9,72.1] | 47.7 [31.6,67.2]† |
| PHE_F | 2.0 | 0.9 [0.2,1.6] | 1.4 [0.6,2.2] | 3.4 [1.8,4.8] | 2.2 [0.9,3.2] | 46.7 [25.8,77.8] | 65.8 [35.8,100.0] |
| PHE_G | 6.0 | 1.0 [0.4,1.7] | 1.5 [0.8,2.3] | 5.1 [3.3,6.8] | 3.5 [2.3,4.5] | 48.4 [28.3,73.1] | 52.4 [31.8,80.9] |

Estimated total body irradiation exposures were calculated without adjustment for FP DCs using ADCI, in accordance with methods presented previously (Li et al. 2019). The FP adjustment was applied when estimating exposure to the irradiated fraction based on Equations 1-3 of the present study. The resulting dose was then used in Equations 4-5 to predicted the fraction of blood irradiated.
\* M denotes usage of a minimal image selection model which in almost all cases selects all images
^ O denotes usage of an optimal image selection model
† values in brackets indicate 95% confidence intervals

Predicted doses of irradiated fractions (Table 3a), predicted fractions of cells irradiated (Table 3b) and *u* values (Table 3c) were determined for all synthetic samples. Although the fraction irradiated estimated from the Dolphin method was generally overestimated in synthetic samples comprised of <50% irradiated cells, the derived values were still consistent with partial body exposures. Tabular data and synthetic sample statistics indicated here refer to specifically synthetic sample set A (sample sets A and B are shown in Supplementary Table 1). For 21 of 24 synthetic samples, dose estimates after optimal image selection were within 1 Gy of the expected predicted exposure of 3 Gy. Twelve of 24 dose estimates were within 0.5 Gy. For CNL, HC, PHE, and DNRI samples, dose-estimated RMSEs were 1.14, 0.64, 0.60, and 0.52 Gy^2^ (corresponding RMSEs of fraction of cells irradiated were 8.06, 26.08, 33.26, 33.28%^2^), respectively. Computed *u* values of synthetic samples are shown in Table 3c. All synthetic samples were correctly predicted to be partially irradiated based on results of the *u* test.

**Table 3a.**
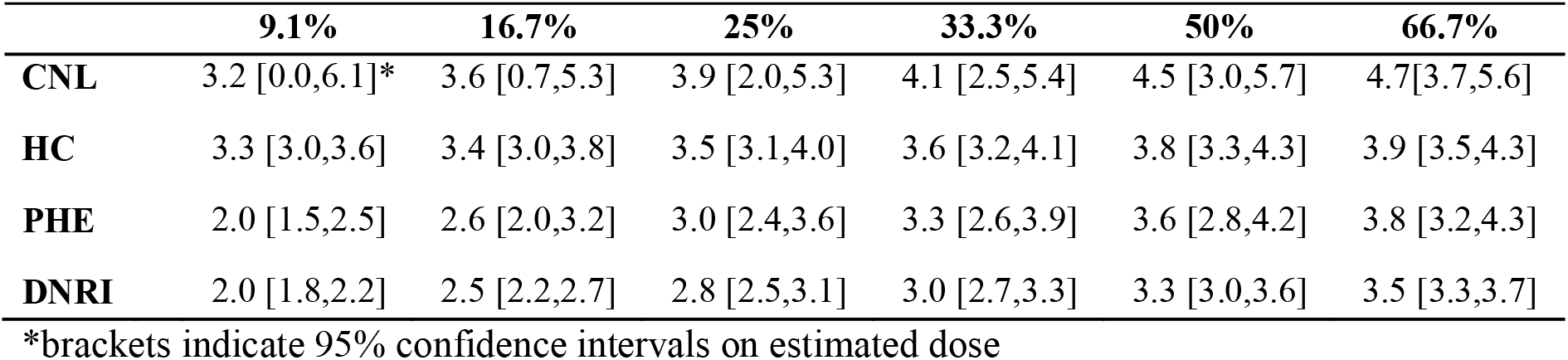
Estimated dose of synthetic, contaminated 3 Gy samples varying the fraction irradiated

**Table 3b.** Estimated irradiated fraction of synthetic, contaminated 3 Gy samples

|  | **9.1%** | **16.7%** | **25%** | **33.3%** | **50%** | **66.7%** |
| --- | --- | --- | --- | --- | --- | --- |
| **CNL** | 1.5[0.2,100.0]* | 9.4<br>[2.3,73.5] | 20.5 [8.0,56.8] | 29.3<br>[13.4,62.5] | 42.8<br>[23.3,72.4] | 52.9<br>[37.4,71.0] |
| **HC** | 39.3 [33.8,45.8] | 47.2<br>[40.0,55.4] | 54.6<br>[46.4,63.7] | 60.9<br>[52.1,70.4] | 71.1<br>[62.0,80.5] | 79.1<br>[72.8,85.4] |
| **PHE** | 51.8 [38.7,70.2] | 55.0<br>[41.4,72.4] | 60.8<br>[46.7,77.5] | 66.4<br>[52.0,82.4] | 76.0<br>[61.8,90.2] | 83.5<br>[74.1,92.4] |
| **DNRI** | 51.6 [44.8,59.8] | 55.3<br>[48.2,63.4] | 60.9<br>[53.6,69.0] | 66.4<br>[59.0,74.3] | 76.0<br>[68.8,83.4] | 83.8<br>[79.0,88.6] |

\*brackets indicate 95% confidence intervals on estimated fraction

**Table 3c.** Computed *u* values of synthetic, contaminated 3 Gy samples for different irradiated fractions

|  | **9.1%** | **16.7%** | **25%** | **33.3%** | **50%** | **66.7%** |
| --- | --- | --- | --- | --- | --- | --- |
| **CNL** | 4.40 | 4.36 | 4.28 | 4.17 | 3.87 | 4.99 |
| **HC** | 11.62 | 8.43 | 6.59 | 5.35 | 3.62 | 3.33 |
| **PHE** | 3.46 | 3.82 | 3.60 | 3.25 | 2.50 | 2.56 |
| **DNRI** | 6.12 | 6.57 | 6.17 | 5.52 | 4.13 | 4.02 |

We also examined potential sources of uncertainty that can contribute to dose estimates obtained with this method. The dose at which 37% of irradiated cells survive, *D_0_* in Eqn. 4, was assigned a value of 3.8 Gy for X-rays in accordance with Barquinero et al. 1997 (Barquinero et al. 1997). However, *D_0_* has previously been reported as 2.7 Gy (Lloyd et al. 1973). We applied the adjusted *D_0_* of 2.7 Gy to HC and DNRI samples (Supplementary Table 1). Varying *D_0_* does not alter the partial body dose estimate but did influence the estimated fraction of cells irradiated. When observing HC and DNRI samples adjusted for FP - 2*SD in synthetic sample set A, mean *F* increased by 7.11% when a *D_0_* of 2.7 Gy was applied. In general, *D_0_* of 3.8 resulted in *F* closer to expected values. A baseline DC rate of 0.00078 was selected based on results from Lloyd et al. (1980). However, the expected baseline DC frequencies range from approximately 0.0005 to 0.002, depending on results from different reports (Lloyd et al. 2006). We created new calibration curves for PHE calibration data at both extreme values of these baseline DC rates. Differences in curve shape and position were negligible, as all adjusted calibration DC frequency values decreased by 0.00028 or increased by 0.00122, when compared with the calibration data presented here. The TP DC count in test samples after FP adjustment is also altered slightly due to the adjustment in baseline DC rate. This adjustment was made according to the number of images in the test sample multiplied by the difference of a new baseline DC rate and our previously applied rate of 0.00078. After application of both extreme baseline DC rates to the calibration curve and test samples, the estimated fraction of cells irradiated for partial body samples was altered by up to 0.98%, with a mean adjustment of 0.38%. Neither of these sources of uncertainty altered the estimated partial body dose.

### Discrimination of partially irradiated samples

The *u* values, dose of irradiated fractions, and fraction of cells irradiated were computed for samples from Interlaboratory exercises of known physical dose from all the laboratories. Whole body dose estimates generated by ADCI were previously published for these exercise samples obtained from HC and CNL (Li et al. 2019). Doses of irradiated fraction and fraction of cells irradiated were calculated for all these exercise samples and synthetic samples before and after FP adjustment, and after both FP adjustment and DC frequency correction (Supplementary Table 1).

To determine whether correction of DC counts for FPs improved discrimination of homogeneously and partially irradiated exercise samples, groups of all exercise and synthetic samples were evaluated separately according to: the *u* test, the level of discrepancy between whole- and partial body estimated exposures, and classification accuracy based on estimated fraction of cells. Samples in which whole body and partial body dose estimates differed by >1 Gy were considered partially irradiated. Samples in which the estimated irradiated fraction of cells was below 75% were defined as partially irradiated. Tables 4a and 4b indicate the numbers of samples correctly classified in each group. Brackets contain 95% confidence intervals of the proportions which were calculated using the modified Wald method (Agresti & Coull 1998). Classification of synthetic partial body samples is significantly improved after FP correction by all these criteria. However, applying FP correction to exercise sample group, which are predominantly comprised of whole-body irradiated samples, reduced classification accuracy. Such corrections are likely counterproductive, since the DC distributions of uncorrected calibration curve samples and exercise samples already compensate for effects of FPs.

**Table 4a.** Percentage of samples appropriately classified as homogeneously or partially irradiated by *u* test, dose discrepancy, or fraction of irradiated cells

| | $u$ value | | Dose discrepancy | | Fraction of irradiated cells | |
| --- | --- | --- | --- | --- | --- | --- |
|  | Exercise<br>n = 25 | Synthetic<br>n = 24 | Exercise<br>n = 25 | Synthetic<br>n = 24 | Exercise<br>n = 25 | Synthetic<br>n = 24 |
| Before FP adjustment | 68<br>[48.3,82.9] | 100<br>[83.7,100] | 56 [37.1,73.4] | 66.7 [46.6,82.2] | 76 [56.3,88.8] | 66.7 [46.6,82.2] |
| FP corrected |  |  | 36 [20.2,55.6] | 91.7 [73,98.8] | 60 [40.7,76.6] | 91.7 [59.1,91.2] |

**Table 4b.** Percentage of samples with ADCI estimated whole body dose >1 Gy appropriately classified as homogeneously or partially irradiated by *u* test, dose discrepancy, or fraction of irradiated cells

| | $u$ value | | Dose discrepancy | | Fraction of irradiated cells | |
| --- | --- | --- | --- | --- | --- | --- |
|  | Exercise<br>n = 20 | Synthetic<br>n = 17 | Exercise<br>n = 20 | Synthetic<br>n = 17 | Exercise<br>n = 20 | Synthetic<br>n = 17 |
| Before FP adjustment | 75<br>[52.8,89.2] | 100<br>[78.4,100] | 60 [38.6,78.2] | 58.8 [36,78.4] | 85 [63.1,95.6] | 52.9 [31,73.8] |
| FP corrected |  |  | 40 [21.8,61.4] | 88.2 [64.4,98] | 75 [52.8,89.2] | 88.2 [46.6,87] |

## Discussion

Partial body exposures to ionizing radiation can be determined consistently and with reasonable accuracy after automated DC identification using unirradiated samples to correct for incorrect DC assignments made by the software. Previously, ADCI analyzed homogeneously exposed calibration and test samples with the same DC detection algorithm, resulting in chromosome misclassifications at similar rates. False positive DCs were anticipated, but their impacts on dose estimation have been masked because they appear at similar rates in all whole body irradiated samples. This self-correcting approach is not feasible for estimating the fraction of cells irradiated of a partially irradiated sample, because this value is determined independently of the corresponding calibration curve. The additional step of estimating and correcting for FPs was required for such samples. We implemented these corrections as a modification of the contaminated Poisson (Dolphin) method (International Atomic Energy Agency 2011) for partial body radiation fraction and dose assessment.

Further improvements will require either identification and removal of additional suboptimal cell images using image selection models or identification of specific FPs in these images. Uniform removal of FPs across the DC distribution, while effective at improving estimated fraction of cells irradiated, does not correct for outlier metaphase cell images with multiple FPs. The FP adjustment method estimates the count of FPs in a set of images but does not identify specific FPs. This is a non-trivial problem, as our previous efforts to target these objects in images resulted in unavoidable loss of true DCs (Liu et al. 2017). Adjustment for FPs influences both the predicted dose and estimated irradiated fraction of a sample. Variance and standard deviation of the dicentric yield of irradiated fraction are increased in samples adjusted for FP due to decreased DC counts. Due to the uniform removal of FPs across the DC distribution, *Y* is unchanged after FP adjustment. Thus, predicted dose differs only due to the adjusted y-intercept of the calibration curve. Ideally, the unirradiated fraction of a partially irradiated test sample and a 0 Gy calibration sample would contain equivalent DC frequency and distribution of DCs in ADCI output. However, in practice this may not be the case. If estimated DC frequency was unusually high in the 0 Gy calibration sample due to FPs detected by ADCI, an excessive number of FP DCs could be removed from test samples resulting in too few DCs remaining in a test sample after FP adjustment. This could result in larger confidence intervals and underestimate the fraction of cells exposed to radiation. Instances of low radiation exposed test samples might be particularly susceptible to this type of overcorrection. Randomized sampling of the unirradiated calibration sample can result in slight differences in the computed fraction of cells exposed (<1%) between different partial body analyses of the same sample. Nevertheless, overall randomized selection of subsets of cells generally corrected dose estimates and fractions consistent with the input partial body composition of the samples.

Of the homogeneously exposed calibration samples from the four laboratories, all samples misclassified by the *u* test were unirradiated (0 Gy), except for the 3 Gy CNL jackknifed test sample. Five of seven exercise samples obtained from PHE were appropriately classified as homogeneously or partially exposed after application of the *u* test. PHE_D, and one partially irradiated sample, PHE_F, were misclassified by the *u* test. PHE_F was exposed to lowest radiation dose and PHE_D was an unirradiated control. The *u* test has been shown to be less reliable for samples with low DC counts (International Atomic Energy Agency 2011).

The u test correctly discriminates whole from partially irradiated samples in all synthetic samples, 75% of exercise samples with whole body dose ≥ 1 Gy and was the best discriminator of the three methods tested. Whole body dose vs partial body dose and estimated fraction of cells irradiated are not as effective at discrimination of whole and partial body irradiation. Samples suspected to be partially irradiated (either because exposure was already known to be inhomogeneous, or from the u test result), FP adjustment of DC counts improved the estimated fraction of cells irradiated in nearly all cases. Removal of samples with estimated whole-body dose < 1 Gy improved correct classification of exercise samples. Most samples in synthetic sample set A were already correctly classified as partially irradiated before removing those with estimated < 1 Gy exposure of the whole-body fraction. Of the synthetic samples with <50% fractional exposure, determination of partially irradiated status is straightforward due to their significant overdispersion of DCs. Only 2 of 24 samples belonging to synthetic sample set B were erroneously classified by the *u* test. Both samples contained 66.7% irradiated cells, suggesting that synthetic samples are more likely to be misclassified by the *u* test as the percentage of irradiated cells approaches 100%.

In some instances, these analyses would have benefited from calibration samples with increased numbers of metaphase cell images, since this constraint limited ADCI’s ability to select high quality cells. Because the pool of metaphase cells available for image selection is halved by the jackknifing process, the pool of high-quality images well suited for automated analysis by ADCI is also halved when examining such samples.

The metaphase cell context is essential in partial body radiation exposure methods that rely on the contaminated Poisson and variant approaches which require the distribution of DCs across all cells (Royba et al. 2019). Partial body dose estimation with ADCI can be completely automated once a calibration curve has been created and the optimal image selection model has been applied. This occurs prior to examination of test samples and is repeated only when a different calibration curve is used to estimate dose. Numerous samples can be classified as either whole- or partially irradiated, with high throughput estimation of low dose exposures and partial body irradiation fractions as low as 9%. Vaurijoux et al. (2012) examined the feasibility of semi-automated DC scoring with DCScore (Metasystems) for partial body exposure determination and recommended manual confirmation of DCs after processing with this software. These authors assessed *in vitro* simulated partial body exposure by Poisson overdispersion in samples consisting of mixtures of 2 Gy irradiated and unirradiated cells, ranging between 5-75% fractions. Six samples had a dose estimated RMSE of 0.62 Gy^2^, compared to the dose estimate RMSE of all samples in synthetic sample set A in the present study of 0.76 Gy^2^. Vaurijoux et al. calculated the estimated fraction of blood irradiated using three different *D_0_* values (2.7, 3.5, 3.8 Gy) and reported the true fraction of blood irradiated fell within the 95% confidence intervals of *F* slightly less than half of the time, across the three *D_0_* values. Similarly, 95% confidence intervals on estimates of *F* calculated by ADCI did not contain the true fraction of blood irradiated in the majority of cases. While the estimate of *F* may currently be unreliable using automated methods presented here, we found that the *u* test reliably detected sample overdispersion when examining simulated partial body exposures in both studies.

Alternative approaches to distinguish whole-from partial-body exposures are expected to exhibit comparable accuracy and dynamic range. ADCI provides unattended analysis and can process sufficient numbers of metaphase cells to achieve accurate exposure levels of the irradiated fraction. Also, the DCA exhibits higher quantitative discrimination and smaller confidence intervals below 6 Gy than other calibrated approaches, eg. PCC, which follow a linear relationship between marker frequency and dose (Lindholm et al. 2010). While fluorescence *in situ* hybridization (FISH) using chromosome paint probes, has been used to detect aberrations, it is not ideally suited for partial body dose estimation due to its comparatively lower aberration detection rates, as only a small subset of chromosomes are examined (Duran et al. 2002). However, the DCA is limited in its ability to accurately assess high dose exposures as cell proliferation is impaired (Sasaki & Norman 1966). Scoring of premature chromosome condensation (PCC) rings has been shown to effectively assess high-dose exposures but has difficulty differentiating whole-from partial-body exposures at low Gy (Romero et al. 2012). Neither the DCA nor PCC ring assay could adequately estimate partial body dose in a simulated triage scenario of 30 dicentrics or 50 metaphase cells; 50 rings or 300 PCC cells (Lindholm et al. 2010). The confidence intervals of whole-body radiation dose estimates of other non-chromosomal assays, including the cytokinesis block micronucleus assay (CBMN), H2AX foci, and protein-based assays are significantly larger than those obtained using the DCA. CBMN did not reliably differentiate partial-from wholebody radiation exposure based on application of the Dolphin and other methods (Mendes et al. 2019). Nevertheless, binary classification of image segmented features from CBMN assay data distinguishes uniformly-from 50%-fractionally irradiated samples (Shuryak et al. 2020). Partial body exposures could also be quantified using next generation sequencing-based RNA-Seq data that distinguishes constitutional-from radiation-specific, alternatively spliced transcript read counts. These features could be incorporated into biochemically inspired-machine learning-based gene expression signatures of ionizing radiation (Dorman et al. 2016; Macaeva et al. 2016; Mucaki et al. 2016; Zhao et al. 2018; Mucaki et al. 2019; Mucaki et al. 2020).

The majority of radiation accident victims, for example, those involved in criticality accidents and inadvertent handling of radioactive materials, receive inhomogeneous exposures (Prasanna et al. 2010). In space, radiation from high energy solar ejecta also produce inhomogeneous dose distributions (Kennedy 2014). In cases of partial body exposure, effective treatment options may differ from those used in homogeneous exposures at the same whole body DC frequency (Prasanna et al. 2010). Partial body dose estimation has been incorporated into the latest version of the ADCI software (https://adciwiki.cytognomix.com/doku.php?id=main:partialbodyestimatedose). Although the dose estimates of partial body exposures generated are not precisely identical to manually determined estimates, the FP elimination method described here was expeditious and produced sufficiently similar dose estimates in both synthetic and actual partially irradiated samples. Large-scale radiation accidents will require both discrimination of homogeneous and partial body exposures as well as timely dose estimation. Complete, integrated automation of the DCA that includes dose estimation will be more expeditious and portable in such accidents when compared with traditional approaches (Rogan et al. 2020).

